# A transformer-based model reveals sparse, stimulus-dependent orientation readout from macaque V1 population activity

**DOI:** 10.64898/2026.09.15.751671

**Authors:** Xin Wang, Shi-Ming Tang, Cong Yu

## Abstract

Orientation information in primary visual cortex (V1) is represented by large populations of neurons with overlapping tuning preferences, yet how this information is selectively read out remains unclear. Here we proposed a transformer-based model to reconstruct oriented Gabor stimuli from two-photon calcium responses of more than 1,000 simultaneously recorded macaque V1 neurons. The model reconstructed stimulus orientation with high precision and revealed, through its self-attention maps, a sparse and stimulus-dependent readout structure. For each stimulus orientation, reconstruction was dominated by a small number of highly weighted readout neurons that were tuned near the presented orientation and showed enhanced effective orientation signals after self-attention modulation. After removal of these neurons and retraining, reconstruction recovered through recruitment of substitute neurons with similar response properties, indicating that sparse readout can be flexibly supported by redundant population encoding. Decoder comparisons showed that a simple linear decoder and a multilayer perceptron recovered orientation less precisely than the transformer, suggesting that the model’s advantage was not explained simply by generic nonlinear decoding capacity. Together, these findings suggest a population-level principle in which redundant orientation encoding supports sparse, stimulus-dependent, and flexible readout.

## Introduction

Orientation selectivity is a fundamental property of neurons in primary visual cortex (V1). Classical single-unit studies established that many V1 neurons respond selectively to stimulus orientation (De Valois et al., 1982; Hubel & Wiesel, 1959, 1962; Ringach et al., 2002; Schiller et al., 1976), and later quantitative work showed that some visual cortical neurons can approach behavioral sensitivity in fine discrimination tasks (Bradley et al., 1987; Parker & Hawken, 1985). These findings raised an important question about visual readout: how is orientation information extracted from neuronal activity? This question becomes especially important at the population level, because a visual stimulus activates hundreds or thousands of V1 neurons with overlapping orientation preferences, diverse response amplitudes, and correlated variability.

Large-scale population activity provides both an opportunity and a challenge for sensory readout. On one hand, redundant population codes can support robust stimulus representations because similar information is available across many neurons (Cunningham & Yu, 2014; Kriegeskorte & Wei, 2021; Stringer et al., 2021; Stringer & Pachitariu, 2024). On the other hand, redundancy creates a readout problem. Accurate decoding may not require all active neurons to contribute equally. Instead, efficient readout may depend on selectively weighting neurons that are most informative under a given stimulus condition. Such neurons cannot be identified solely from their intrinsic orientation tuning, because their effective contribution also depends on response amplitude, variability, shared noise, population context, and the form of the readout itself (Averbeck et al., 2006; Butts & Goldman, 2006; Kang et al., 2004; Series et al., 2004).

Current neural decoders have provided powerful tools for quantifying the information contained in neuronal populations. Linear models recover stimulus information through fixed weighted pooling, whereas nonlinear models such as multilayer perceptrons (MLPs) can capture more complex input–output relationships (Hornik, 1991; Hornik et al., 1989). Such nonlinear models are widely used in neural decoding to capture relationships between population activity and stimulus or behavioral variables that may not be well described by linear mappings (Glaser et al., 2020; Livezey & Glaser, 2021; Mathis et al., 2024). Although their learned parameters remain fixed after training, nonlinear feedforward models can still exhibit input-dependent sensitivity. Therefore, they do not provide an explicit, condition-specific weighting matrix showing how individual neuronal contributions vary across stimulus conditions, limiting their usefulness for directly examining stimulus-dependent population readout.

A model that combines accurate decoding with stimulus-dependent readout interpretation is therefore needed. Transformer architectures provide a framework through self-attention mechanisms, which compute relationships among input neurons based on the current activity (Vaswani et al., 2017). Unlike conventional fully-connected decoders with fixed weights, self-attention generates stimulus-dependent weighting among neuronal representations, providing a computational approach for examining how information from different neurons is selectively weighted during stimulus reconstruction. Importantly, we do not interpret self-attention as a direct biological mechanism, but rather use it as a computational tool to investigate possible principles of population readout.

Here, we developed a transformer-based model to reconstruct oriented Gabor stimuli from two-photon calcium responses of more than 1,000 simultaneously recorded macaque V1 neurons. Compared with conventional linear and nonlinear fully-connected feedforward decoders, the transformer achieved substantially higher orientation reconstruction accuracy.

We further analyzed stimulus-dependent attention patterns to examine the organization of neuronal contributions during reconstruction. These analyses revealed that neurons receiving high attention weights exhibited biologically meaningful properties, indicating that the model-derived readout structure was associated with functionally relevant aspects of cortical population organization.

## Results

Neuronal data used for model training and testing were obtained from seven awake, fixating macaques, each contributing one imaging field of view (FOV). Each FOV contained responses from 1,039 to 1,903 simultaneously recorded superficial-layer V1 neurons (150 μm deep from the cortical surface) measured with two-photon calcium imaging (Fig. 1A-C). The FOV size was 850 x 850 mm^2^, approximately corresponding to one orientation hypercolumn (Hubel & Wiesel, 1977).

**Figure 1.**
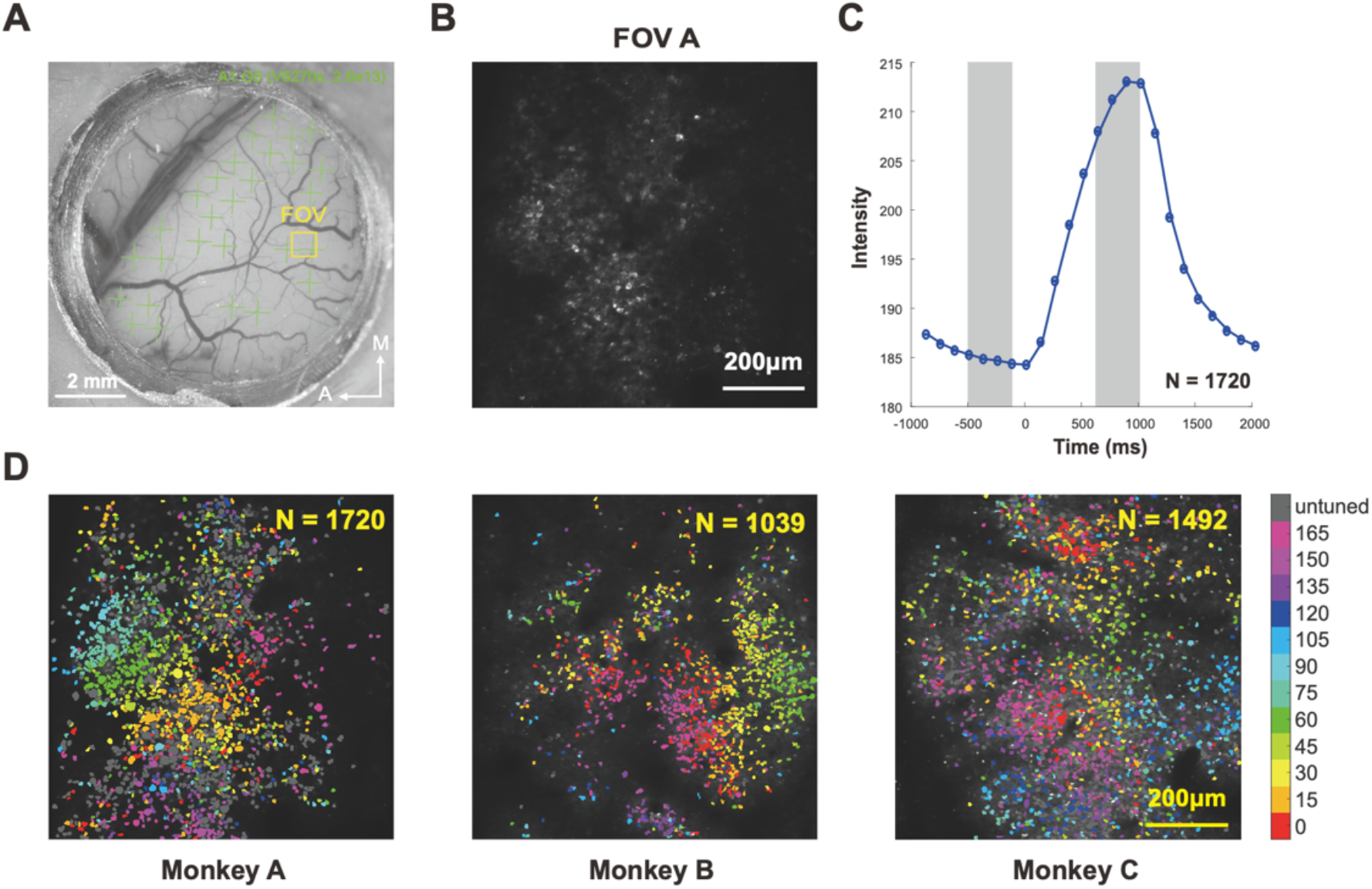
Two-photon calcium imaging and orientation functional maps. **A.**Exemplar vascular maps of Monkey A. Green “+” signs indicate the viral injection regions. The yellow box on the vascular map indicates the FOV chosen for imaging (850 x 850 μm^2^). **B**. The average two-photon image of Monkey A over a session. **C**. The time course of calcium responses of Monkey A. The curve is averaged over all neurons. Each dot indicates the mean response intensity calculated from one frame (8 fps after averaging every 4 frames). Error bars represent ±1 SEM. Shaded areas before or after the stimulus onset denote the 4 frames that were used for calculating the baseline (*F0)* and response (*F)* values, respectively. **D**. Example cellular orientation functional maps from Monkeys A, B, and C. Each dot indicates one individual neuron and the color indicates the neuron’s orientation preference. The grey color indicates neurons with no significant orientation tuning (p > 0.01, Friedman test). 82.8% of the identified neurons were found to be orientation selective over seven FOVs.

### Transformer-based decoder enables high-precision reconstruction

The transformer-based decoding model comprised five single-layer components: embedding, positional encoding, self-attention, unembedding, and a fully-connected output layer (Fig. 2A). For each stimulus condition, the input consisted of a condition-averaged population response vector, and the output was the reconstructed Gabor image.

**Figure 2.**
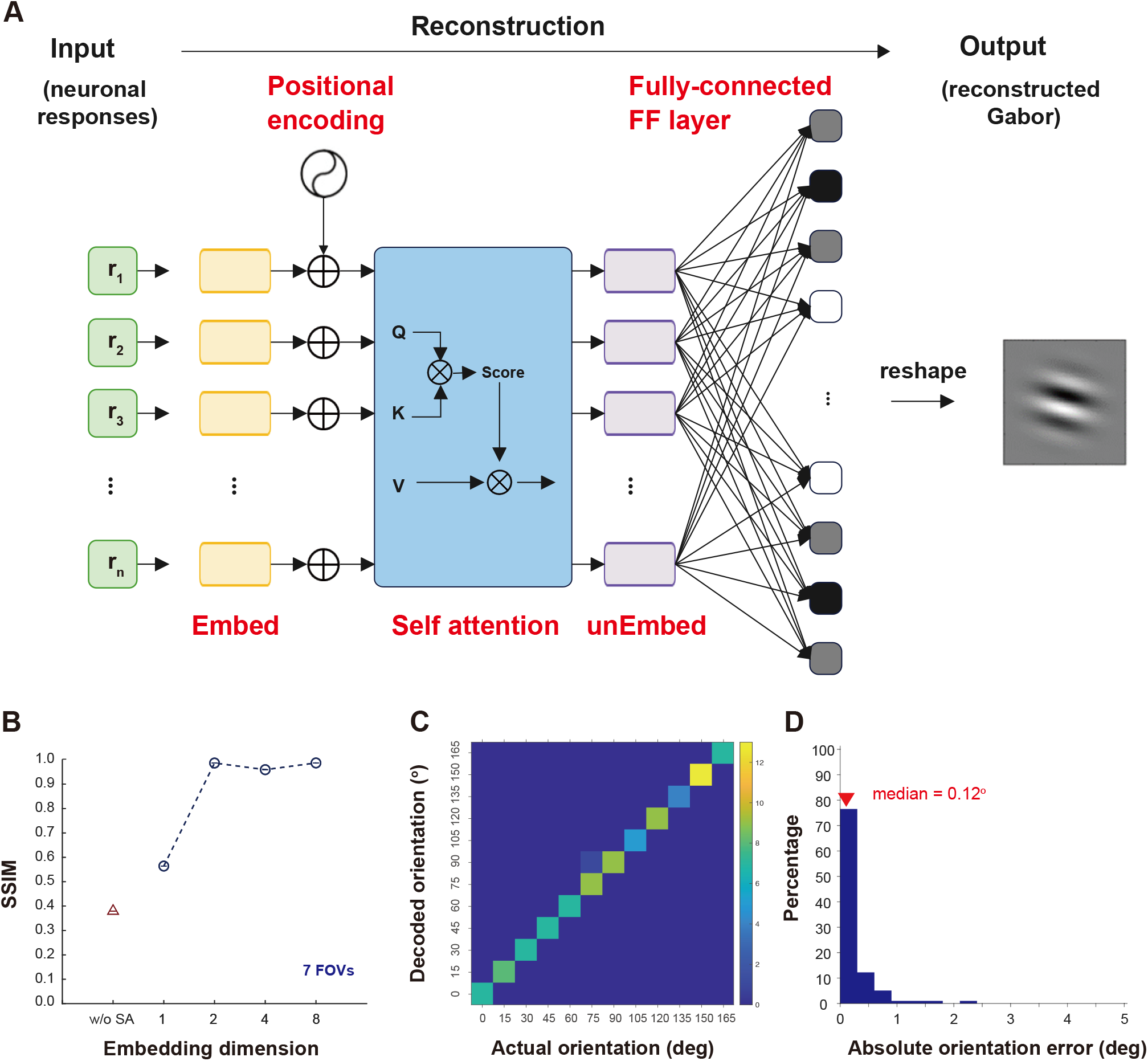
The transformer-based model. **A.**The model architecture. Details are described in the text. **B**. The model performance under various embedding dimensions. To determine the appropriate dimensionality within the embedding module, we calculated SSIM in the training phase when the number of dimensions varied at 1, 2, 4, and 8. Mean SSIM exceeded 0.95 when the embedding dimensionality was ≥2. w/o SA: without self-attention. Thus, we chose a model with two embedding dimensions. **C**. Decoded orientations versus actual stimulus orientations in test trials. **D**. Distribution of absolute orientation errors.

The embedding module mapped neuronal responses onto a two-dimensional latent space (Fig. 2B) using neuron-specific learnable weights, enabling the latent representation to preserve the distinct properties of individual neurons. A fixed two-dimensional positional encoding then arranged neurons according to the 0°-180° orientation cycle, such that neurons with similar preferred orientations were positioned nearby in the input vector, reflecting orientation clustering observed in V1 (Hubel & Wiesel, 1962). The core of the model was a single-layer self-attention module, which computed how each neuron’s representation was weighted by the representations of other neurons through Query, Key, and Value operations(Vaswani et al., 2017). In the present context, self-attention provides a decoding-level measure of how each neuron’s contribution depends on others during stimulus reconstruction. The resulting two-dimensional representations were then compressed into a one-dimensional vector through unembedding and mapped to image pixels by a fully-connected output layer, yielding the reconstructed Gabor image.

The model was trained for image reconstruction and achieved high performance, with a mean structural similarity index (SSIM) of 98.3%. We then evaluated orientation decoding performance by extracting the orientation of each reconstructed image (see Methods). The decoded orientations closely matched the true stimulus orientations (Fig. 2C), with a median absolute error of 0.12° (Fig. 2D). For all analyses below, we focused on 12 orientations per FOV at the spatial frequency and stimulus size preferred by most neurons in that FOV.

We next asked whether the transformer’s high orientation precision could be achieved by simpler feedforward decoders. A direct linear decoder and a nonlinear multilayer perceptron (MLP), both trained to predict orientation from the population response (see Methods), produced median orientation errors of 2.83° and 2.92°, respectively, compared with 0.12° for the transformer. Because these decoders differed from the transformer in both output representation and training objective, we also examined a linear image-reconstruction control using the same Gabor targets and reconstruction objective. This control is functionally equivalent to removing the embedding and self-attention modules from the transformer while retaining a linear mapping from population responses to the reconstructed image. Its removal reduced SSIM (Fig. 2B) and yielded a median orientation error of 2.95°, again substantially larger than that of the transformer. Thus, the transformer’s higher precision cannot be explained simply by differences in output representation or training objective, although this comparison does not isolate self-attention from the other architectural components.

Beyond improved decoding accuracy, the transformer provides an additional advantage over conventional fully-connected decoders: its stimulus-dependent attention patterns characterize the structure of the model’s readout weighting. Unlike fixed-weight feedforward decoders, in which neuronal weights remain invariant across stimulus conditions, the transformer dynamically assigns stimulus-dependent self-attention weights among neurons.

This provides a computationally interpretable framework for examining how population activity is selectively weighted during orientation reconstruction.

### Self-attention reveals sparse, stimulus-dependent orientation readout

An attention score, which ranges from 0 to 1, quantifies how strongly the decoder weights one neuron’s Value representation when computing another neuron’s representation during reconstruction. A score closer to 1 indicates a stronger contribution, while a score closer to 0 indicates little or no contribution. Based on these scores, the model identifies which neuronal representations receive the strongest model-assigned weights during orientation reconstruction.

The attention scores received by neurons in response to an exemplar 0° stimulus, together with their means, are shown in Figs. 3A–B for exemplar FOVs A–C. Only a small number of neurons received strong attention scores from the rest of the population during decoding. We refer to these neurons operationally as highly weighted readout neurons under a given stimulus condition. To identify these neurons quantitatively (Fig. 3C), we first ranked neurons within each FOV by the mean attention they received from all other neurons. We then thresholded the self-attention matrix, before multiplication with the Value component, by progressively lowering the attention-score cutoff (equivalently, by progressively increasing the top percentage of attention scores retained). In the attention-retained condition, only attention scores above the cutoff were retained, whereas in the attention-ablated condition, those same supra-threshold scores were set to zero. For each cutoff, we quantified the mean difference between the reconstructed and true stimulus orientations. The effective threshold was defined as the highest attention-score cutoff—or equivalently, the smallest retained fraction of attention scores—at which the mean absolute orientation error was no greater than 5°, and attention scores above this cutoff were defined as effective attention scores. Highly weighted readout neurons were then defined as the neurons receiving these effective attention scores.

**Figure 3.**
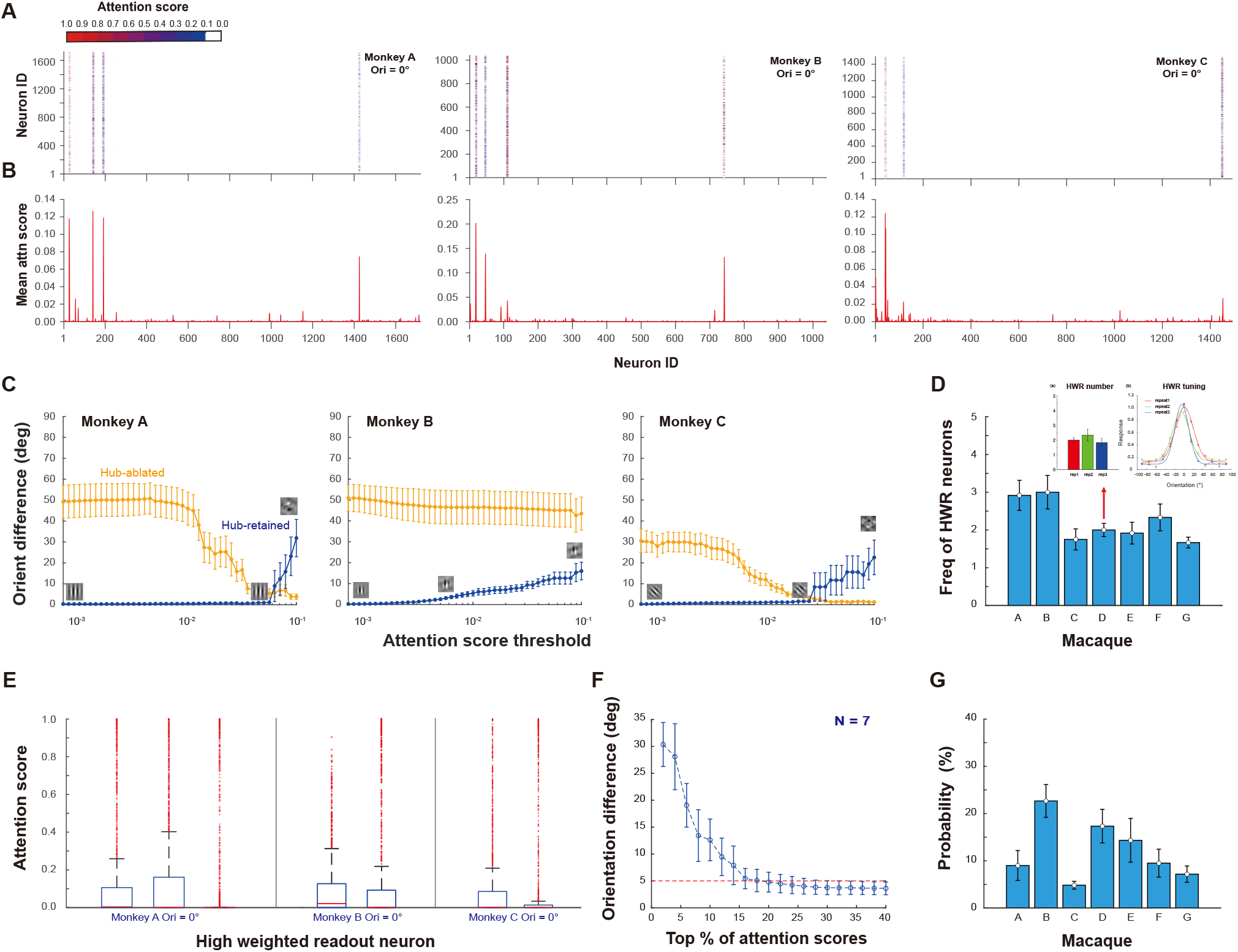
Self-attention maps and highly weighted readout neurons. **A.**Attention score matrices from three exemplar FOVs, showing pairwise attention scores between neurons for an exemplar 0° stimulus. **B**. Mean attention score received by each neuron from all neurons in the attention maps shown in **A** (i.e., column means). **C**. Mean differences between the reconstructed orientation and the true stimulus orientation in the attention-retained and attention-ablated conditions. In both conditions, thresholding was applied to the attention map before multiplication with the Value term. In the attention-retained condition, only attention scores above the threshold were retained, while all scores below the threshold were set to zero. In the attention-ablated condition, attention scores above the threshold were set to zero, whereas all remaining scores were retained. As the threshold was progressively lowered, more high-attention-score neurons were included in the attention-retained condition but removed in the attention-ablated condition. The 5° criterion refers to a mean orientation difference of 5° or less between the reconstructed and true stimulus orientations. Error bars indicate ±1 SEM. **D**. Mean frequencies of highly weighted readout neurons across 12 orientations for each FOV. Middle: mean number of highly weighted readout neurons obtained in three cross-validation repeats, each run with an independent random seed. Right: average orientation-tuning curves of highly weighted readout neurons from the three repeats. **E**. Box plots of the distributions of attention scores from all neurons to highly weighted readout neurons in three exemplar FOVs. Because the distributions are highly skewed and the medians are near zero, only the upper quartile, upper whisker, and outliers above the upper whisker are clearly visible. **F**. Mean orientation difference in reconstructed stimulus as a function of the top percentage of attention scores retained. The red dashed line indicates the 5° criterion. **G**. Probability that highly weighted readout neurons received effective attention scores for each FOV.

We found that only 2-3 highly weighted readout neurons on average were sufficient across the 12 reconstructed orientations (Fig. 3D, left). Thus, accurate orientation decoding was dominated by a very small subset of neurons rather than the full population. The number of identified highly weighted readout neurons was stable at the population level across model initializations with different training and testing splits and random seeds. For example, in Monkey D, three independent runs yielded similar numbers of highly weighted readout neurons, and although the exact neurons varied, their tuning functions remained highly consistent (Fig. 3D, middle and right). The attention scores received by highly weighted readout neurons were highly skewed (Fig. 3E). Among the incoming attention scores received by the identified neurons, the strongest 12.12% were sufficient to reach criterion (Figs. 3F-G). Because only a few recipient neurons were involved, these entries represented 0.021% of the full attention matrix. This highly skewed readout structure is consistent with the idea that a small minority of strongly weighted signals can support most immediate decoding demands (Buzsáki & Mizuseki, 2014). Together, these results indicate that the model implements a highly concentrated effective readout structure, suggesting a possible computational principle for selective neural readout.

### Highly weighted readout neurons have biologically meaningful response properties

Although identified through a data-driven decoder, highly weighted readout neurons exhibited biologically meaningful response properties, indicating that they were not assigned arbitrarily across the recorded population. First, the preferred orientations of highly weighted readout neurons (shown in red) closely matched the actual stimulus orientations (0° in the current example for three exemplar FOVs, Fig. 4A). The median difference over 12 orientations was approximately 5-10° across all FOVs, suggesting that highly weighted readout neurons were preferentially drawn from neurons tuned near the presented orientation (Fig. 4A). Second, the mean attention scores of highly weighted readout neurons were negatively correlated with their mean noise correlations to other neurons within the same FOVs when data from six FOVs were pooled (r = -0.267, p < 0.001; Fig. 4B). This indicates that more dominant highly weighted readout neurons (i.e., those with higher attention scores) tended to exhibit lower shared variability, making their activity more independent and potentially more effective for orientation decoding (Series et al., 2004). Third, highly weighted readout neurons tend to be located in regions with above-average but not extreme orientation gradients. We divided the mean distribution of normalized orientation gradients at highly weighted readout neuron locations (Fig. 4C bottom-left) by that over all pixels across all FOVs (Fig. 4C bottom-middle), which revealed that highly weighted readout neurons were overrepresented in regions with orientation gradients of 0.6-0.8 (Fig. 4C bottom-right). The apparent peak at 0.9-1.0 (gray bar) likely reflected the sparsity of high-gradient pixels in these FOVs, which inflated the ratio. This distribution suggests that highly weighted readout neurons were preferentially located in regions with intermediate local variation in orientation preference, although the functional significance of this spatial bias remains uncertain.

**Figure 4.**
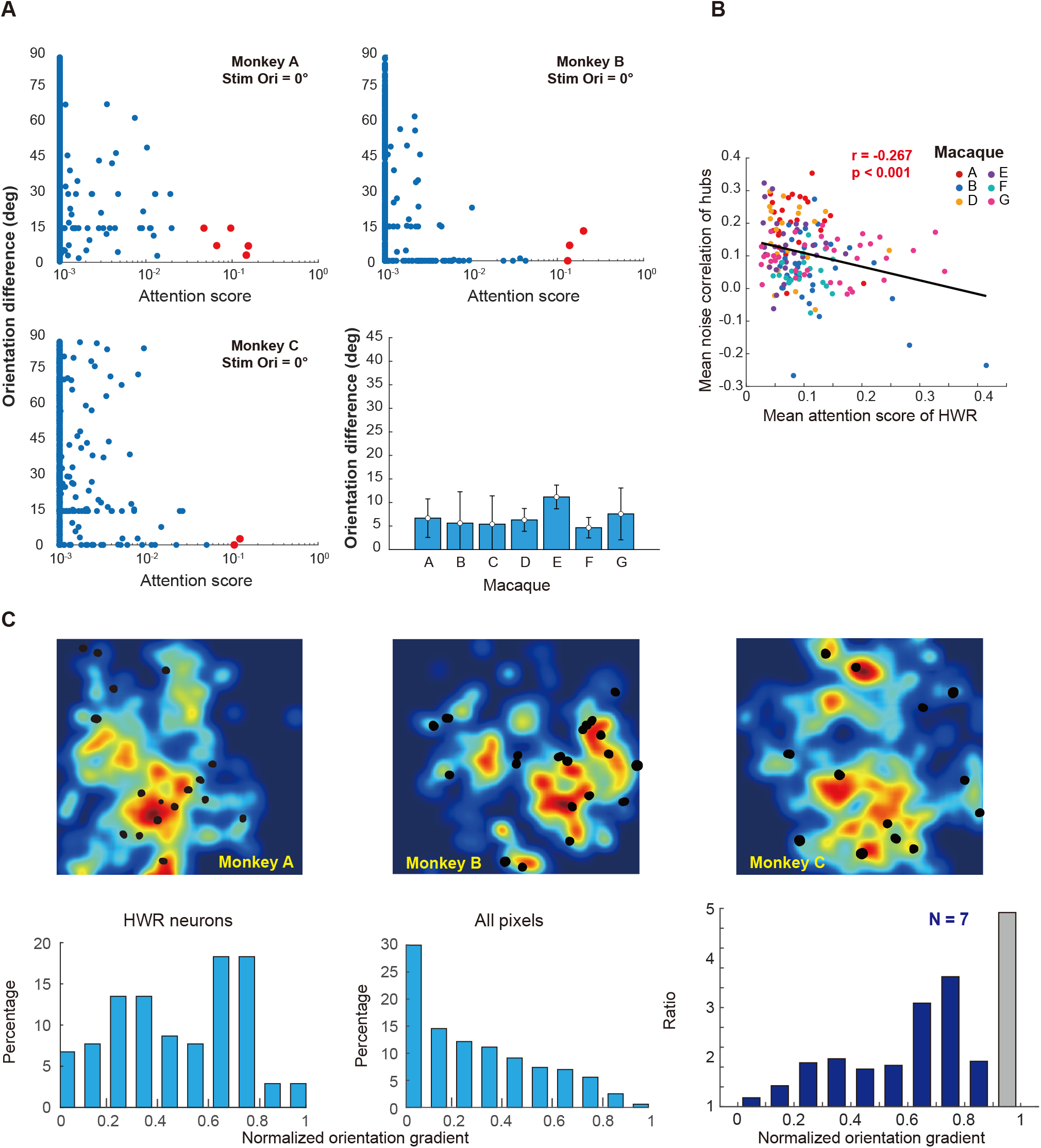
Biological relevance of highly weighted readout neurons. **A.**Differences between the preferred orientations of highly weighted readout neurons and the stimulus orientation in three exemplar FOVs and the histogram plot of median difference between the preferred orientations of highly weighted readout neurons and the actual stimulus orientations across 12 stimulus orientations for each FOV. **B**. Negative correlation between highly weighted readout neurons’ attention scores and their mean noise correlations to other neurons. Each dot represents a highly weighted readout neuron. Colors indicate different FOVs. The black line shows the linear fit of all data points. Data were pooled from six FOVs because Macaque C was excluded due to laser instability precluding reliable estimation of noise correlations. **C**. Spatial distribution of highly weighted readout neurons (black dots) overlaid on orientation gradient maps from three exemplar FOVs. Bottom-left and bottom-middle panels: Histograms show the raw distributions of normalized orientation gradients for highly weighted readout neurons (left) and all pixels (middle). Bottom-right panel: Ratio of the distribution of normalized orientation gradients at highly weighted readout neuron locations to the pixel-level orientation gradient distribution. Data were pooled from all seven FOVs.

### Attention weighting sharpens and amplifies the effective orientation profiles of highly weighted neurons

Highly weighted readout neurons were among the most responsive neurons in each FOV, ranking within the top 1.08% ± 0.35% of the population in response amplitude. We asked how self-attention reshaped population responses to support orientation readout. To address this, we compared the mean orientation tuning functions of highly weighted readout neurons and the top 1% ranked response-matched comparison neurons after aligning each neuron’s tuning curve to its preferred orientation. Self-attention reweighted neuronal responses across stimulus orientations according to their attention scores (Fig. 5A).

**Figure 5.**
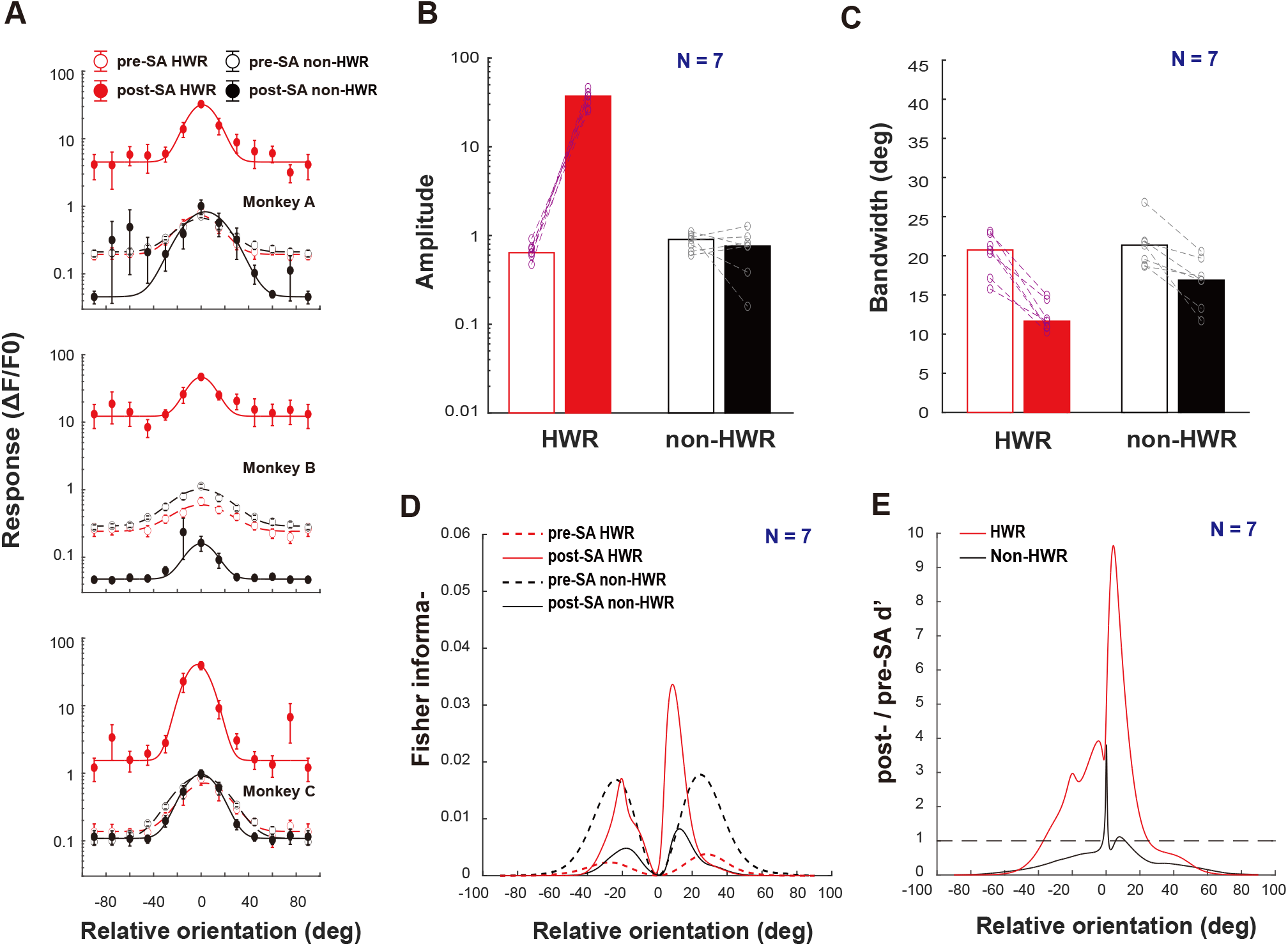
Self-attention modulation of orientation information from highly weighted readout neurons and response-matched comparison neurons. **A.** Exemplar mean orientation tuning functions from three FOVs for highly weighted readout neurons and response-matched comparison neurons before and after self-attention modulation. Note that the y-axis is on a log scale. SA - self-attention. **B-C**. Amplitude and bandwidth of the mean orientation tuning functions of highly weighted readout neurons and response-matched comparison neurons before and after self-attention modulation. For each FOV, these quantities were computed from the neuron-averaged tuning functions in A and then summarized across 7 FOVs. **D**. Fisher information for highly weighted readout neurons and response-matched comparison neurons before and after self-attention modulation as a function of relative orientation. **E**. Post-/pre-self-attention d’ ratios for highly weighted readout neurons and response-matched comparison neurons as a function of relative orientation. The d’ ratio was computed as the square root of the ratio of post-to pre-SA Fisher information. Curves show means across 7 FOVs.

Before self-attention modulation, the mean orientation functions of highly weighted readout neurons and similarly ranked response-matched comparison neurons displayed no significant differences in amplitude (p = 0.053; independent samples t-test) and bandwidth (p = 0.488). However, after self-attention modulation, the mean orientation tuning function of highly weighted readout neurons showed substantial increases in amplitude (p < 0.001; Fig. 5B), as well as a significant decrease in bandwidth from 20.19° ± 1.05° to 12.15° ± 0.69° (p < 0.001; Fig. 5C). In contrast, the mean orientation tuning function of response-matched comparison neurons showed no significant changes in amplitude (p = 0.458; Fig. 5B), and a lesser but significant decrease in bandwidth from 21.26° ± 1.06° to 16.63° ± 1.21° (p = 0.006; Fig. 5C). The post-self-attention orientation tuning functions of highly weighted readout neurons and response-matched comparison neurons differed significantly in amplitude (p < 0.001) and bandwidth (p = 0.007). These results indicate that, within the model, self-attention reweighted population responses by amplifying signals from highly weighted readout neurons while reducing the relative contribution of response-matched comparison neurons, thereby improving orientation readout.

We calculated the Fisher information (Averbeck et al., 2006) for both highly weighted readout neurons and response-matched comparison neurons before and after SA modulation, for which we did not consider noise correlation, because the responses were averaged over 12 trials for each condition. Before self-attention modulation, highly weighted readout neurons exhibited a localized Fisher information peak around 20° from the stimulus orientation (Fig. 5D). After self-attention modulation, this peak shifted closer to 10° and became substantially more pronounced, indicating an enhanced orientation discrimination ability. The enhancement is further reflected in Fig. 5E, where the d’ ratio, which is the square root of the ratio of post-to pre-self-attention Fisher information, revealed that self-attention modulation of responses from highly weighted readout neurons significantly increased orientation sensitivity, with ratios exceeding 1 across a wide range of orientations and peaking at 4.77°.

In contrast, after self-attention modulation, the Fisher information from response-matched comparison neurons significantly decreased, except for a small range of relative orientation from 0° to approximately 15° where the Fisher information remained unchanged (Fig. 5D). This indicates reduced orientation sensitivity, particularly at larger relative orientations. Specifically, the d’ ratios for response-matched comparison neurons were mostly below 1, with one exception where the ratios were near or slightly above 1 around 15° relative orientation (Fig. 5E). A further exception occurred for a very narrow range around 0°, where the d’ ratios were substantially above 1. This was likely a spurious effect due to the extremely low Fisher information before self-attention modulation (Fig. 5D). Together, these results show that self-attention selectively amplifies the most informative signals from highly weighted readout neurons while suppressing signals from other neurons, thus enabling efficient readout of core orientation information.

### Redundant V1 population activity supports substitute sparse readout after neuron removal

The identified highly weighted readout neurons were not indispensable in population orientation coding. We performed an ablation experiment in which the identified highly weighted readout neurons were removed and the model was retrained. The retrained model could reconstruct the stimuli equally well with a mean SSIM of 98.9%. This is because the model recruited substitute neurons whose amplitudes were significantly enhanced (p < 0.001), while their tuning bandwidths were significantly reduced (p = 0.009 excluding outlier Monkey F), under self-attention (Fig. 6A–C). The number of substitute neurons did not differ significantly from that of the original highly weighted readout neurons (p = 0.204 excluding outlier Monkey B, Fig. 6D). They were functionally similar to the original neurons as they were also tuned to the stimulus orientation (Fig. 6E). Although the original response amplitudes of substitute neurons were significantly reduced compared to those of highly weighted readout neurons (p = 0.015, Fig. 6E), the probability of substitute neurons receiving effective attention scores from other neurons to achieve the criterion orientation difference (5°) was significantly increased (p < 0.001; Fig. 6F) to compensate.

**Figure 6.**
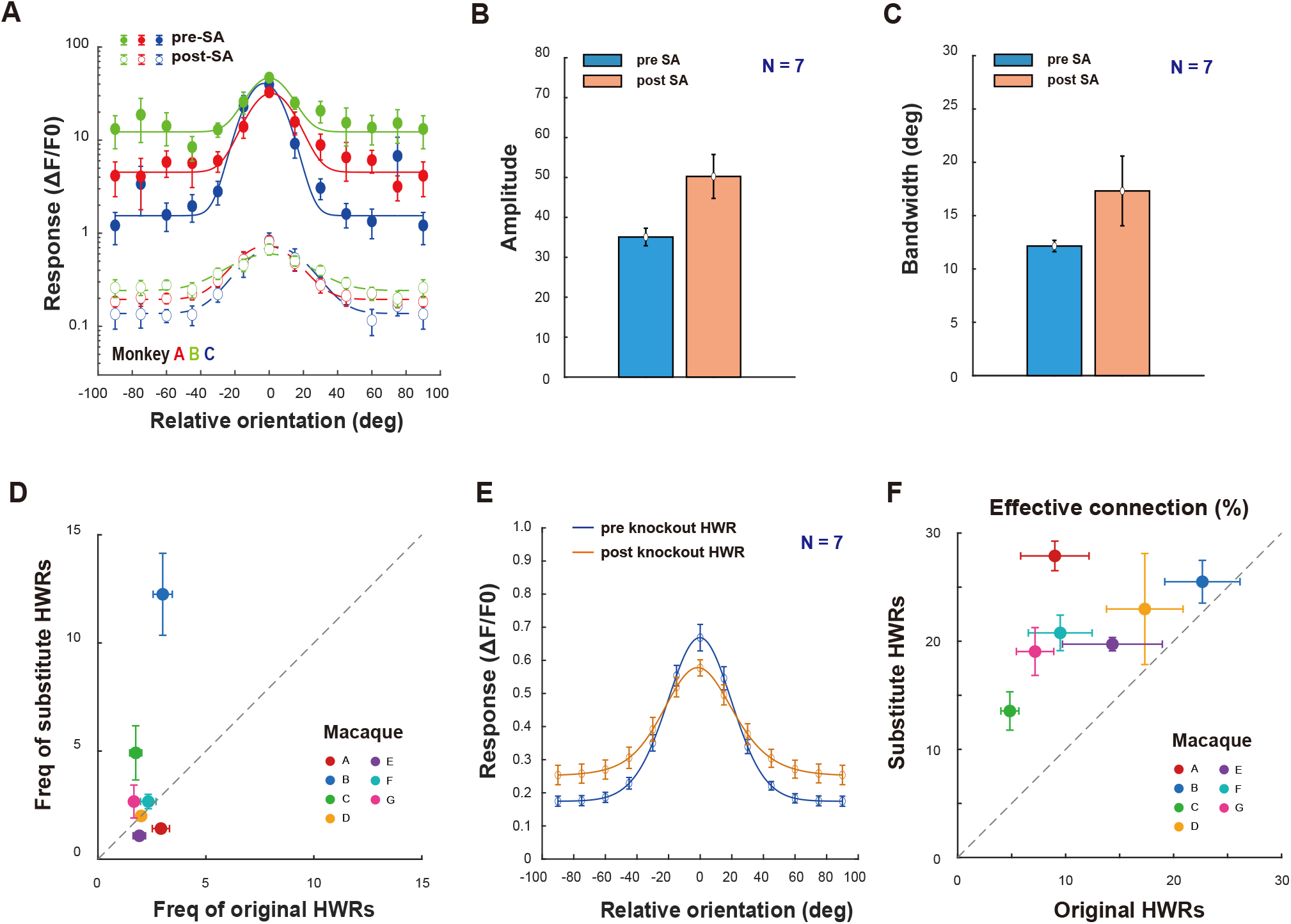
Effects of ablating highly weighted readout neurons and subsequent model retraining. A. Exemplar mean orientation tuning functions of substitute neurons before and after self-attention modulation. B-C. Amplitude and bandwidth of orientation tuning functions of substitute neurons before and after self-attention modulation. D. The mean frequencies of original highly weighted readout neurons vs. substitute neurons. E. Mean orientation tuning functions of original highly weighted readout neurons and substitute neurons over all FOVs before self-attention modulation. F. Mean probabilities of original highly weighted readout neurons vs. substitute neurons receiving effective attention scores. Error bars indicate ±1 SEM.

## Discussion

We found that orientation information in macaque V1 is redundantly represented across large neuronal populations but can be read out through a sparse, stimulus-dependent structure revealed by a transformer-based model. The model reconstructed oriented Gabor stimuli with high precision, providing an empirical benchmark for the orientation information available in the recorded population. Its self-attention maps showed that reconstruction for each orientation was dominated by a small number of highly weighted readout neurons that were tuned near the presented orientation, tended to show lower shared variability, and exhibited stronger and sharper effective orientation signals following self-attention. When these neurons were removed, reconstruction recovered through recruitment of substitute neurons with similar response properties. These findings suggest that redundant orientation representations in macaque V1 can support precise, flexible, and sparse readout.

These results link redundant population coding with sparse effective readout. Orientation information in V1 is distributed across many neurons with overlapping tuning preferences (De Valois et al., 1982; Hubel & Wiesel, 1959, 1962; Ringach et al., 2002; Schiller et al., 1976) . Such redundancy can improve robustness by making similar stimulus information available across multiple neurons (Cunningham & Yu, 2014; Kriegeskorte & Wei, 2021; Stringer et al., 2021; Stringer & Pachitariu, 2024). However, accurate readout does not require all informative neurons to contribute equally. In our model, a large redundant population provided the substrate for reconstruction, but the effective readout was concentrated on a small number of highly weighted neurons. Thus, sparse readout does not imply sparse representation. Instead, sparse and stimulus-dependent readout can be built on top of a redundant population code, consistent with proposals that small subsets of neurons can dominate effective sensory readout under specific conditions (Barlow, 1972; Ince et al., 2013; Kafashan et al., 2021; Yoshida & Ohki, 2020).

The highly weighted readout neurons should not be interpreted as a fixed anatomical or physiological neuron class. They were defined operationally by the model as neurons receiving strong effective attention during stimulus reconstruction. Their identity changed across stimulus orientations, and substitute neurons emerged after removal of the original set. Thus, “highly weighted” status reflects a flexible readout role rather than an intrinsic property of a neuron. Nevertheless, these neurons showed biologically meaningful response properties: they were tuned near the presented orientation, showed relatively low shared variability, and were located in regions with above-average but not extreme orientation gradients. These properties are consistent with population-coding work showing that readout-relevant information depends not only on tuning, but also on response amplitude, variability, correlations, and population context (Averbeck et al., 2006; Butts & Goldman, 2006; Kang et al., 2004; Series et al., 2004).

Self-attention did more than select neurons. Before self-attention modulation, highly weighted readout neurons were not simply distinguished by unusually narrow intrinsic tuning. After modulation, their effective tuning functions became stronger and sharper, and their Fisher information increased near the stimulus orientation. Thus, self-attention implemented a model-derived readout operation that amplified and sharpened the signals most useful for the current reconstruction while reducing the relative contribution of less useful signals. The ablation analysis further showed that these neurons were not uniquely indispensable: when they were removed, the model recruited substitute neurons with similar functional properties. This indicates that the V1 population contains multiple potential sparse routes for recovering the same orientation information.

The decoder comparisons clarify the computational role of the transformer. It provided the most precise orientation recovery among the tested architectures, with a median error of 0.12°, compared with approximately 2.8–2.9° for the linear decoder and MLP. Because additional variability is likely to arise during downstream processing and decision formation, the feedforward decoders may leave insufficient margin for supporting fine orientation discrimination. The transformer was also uniquely informative because its explicit self-attention matrix revealed that high-precision reconstruction was supported by sparse, stimulus-dependent weighting of a small number of highly weighted readout neurons. Thus, the transformer served not only as an accurate decoder, but also as an interpretable model for exposing the readout structure underlying orientation reconstruction from V1 population activity.

More broadly, attention-based models may provide a useful framework for investigating how visual information is represented and selectively read out from large neural populations. Recent studies have used transformer architectures to model dynamic routing of retinotopic visual features to category-selective human cortex (Adeli et al., 2026) and to predict and interpret responses in higher visual cortex across subjects and stimuli using in-context learning (Yu et al., 2026). Related transformer-based architectures in our previous studies captured orientation coding under continuous flash suppression in macaque V1 (Chen et al., 2026) and orientation serial dependence in macaque V1/V2 using cross-attention (Wang et al., 2026). Together with the present results, these studies suggest that attention-based models provide a flexible computational framework for studying dynamic population readout across different visual computations.

## Materials and Methods

### Neuronal data

The V1 neuronal data used in the current analysis were obtained from seven awake, fixating macaques (all males, 4-6 years old), one FOV per animal. Imaging was performed using two-photon calcium imaging over cortical areas of 850 x 850 μm^2^. The dataset includes previously published ones from Monkeys D-G(Ju et al., 2022; Ju et al., 2021), as well as unpublished ones from Monkeys A–C. All recordings were performed at a cortical depth of 150 µm.

For each FOV, neuronal responses to a Gabor stimulus were recorded under 216 unique stimulus conditions, defined by a full factorial combination of 12 equally spaced orientations, 6 spatial frequencies in 1-octave steps from 0.25 to 8 cpd, and 3 stimulus sizes in wavelength units. Each stimulus condition was presented 12 times, and the final response of each neuron was defined as the average across repeats to reduce trial-level variability and improve the signal-to-noise ratio.

Then fluorescence changes were associated with corresponding visual stimuli through the recorded time sequence information. By subtracting the mean of the four frames before stimulus onset (F0) from the average of the 6th–9th frames after stimulus onset (F) across five or six repeated trials for the same stimulus condition, the differential image (ΔF = F – F0) was obtained.

For a specific FOV, imaging data were aligned using a normalized cross-correlation-based translation algorithm, with reference images selected from recordings showing the least head movement and highest self-correlation. Then the regions of interest (ROIs) or candidate cell bodies were determined through sequential analysis of 216 differential images in the order of spatial frequency (6), size (3), and orientation (12) (6 × 3 × 12 = 216). The first differential image was filtered with a band-pass Gaussian filter (size = 2 to 10 pixels), and connected subsets of pixels (>25 pixels, which excluded smaller vertical neuropils) with average pixel value > 3 SD of the mean brightness were selected as ROIs. Then, the areas of these ROIs were set to mean brightness in the next differential image before the bandpass filtering and thresholding were performed. This measure gradually reduced the standard deviations of differential images and facilitated the detection of neurons with relatively low fluorescence responses. If a new ROI and an existing ROI from the previous differential image overlapped, the new ROI was retained as a separate ROI if the overlapping area (OA) < 1/4 ROI_new_, discarded if 1/4 ROI_new_ < OA < 3/4 ROI_new_, and merged with the existing ROI if OA > 3/4 ROI_new_. Merging helped smooth the contours of the final ROIs. This process was repeated for all 216 differential images twice to select ROIs. Finally, the roundness for each ROI was calculated as:

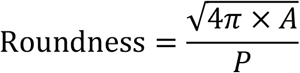

where A was the ROI’s area and P was the perimeter. Only ROIs with roundness larger than 0.9, which would exclude horizontal neuropils, were selected for further analysis. The sizes of the ROIs ranged from 3.3 to 19.3 μm in diameter, with an average of 8.9 ± 1.3 μm.

The ratio of fluorescence change (ΔF/F0) was then calculated as a neuron’s response to a specific stimulus condition. For a specific neuron’s response to a specific stimulus condition, the F0_n_ of the n-th trial was the average of 4 frames before stimulus onset (–500 to 0 ms), and F_n_ was the average of the fifth to eighth frames after stimulus onset (500 to 1,000 ms). F0_n_ was then averaged across 12 trials to obtain the baseline F0 for all 12 trials (to reduce noise in the response calculations), and ΔF_n_/F0 = (F_n_ – F0)/F0 was taken as the neuron’s response to this stimulus at the n-th trial.

Several steps were then taken to determine whether a neuron was tuned to orientation. For each condition, the orientation, spatial frequency, and size (σ) that produced the maximal response among all conditions were selected. Then, responses to the other 11 orientations were determined at the selected spatial frequency and size. Second, to identify orientation-tuned neurons, a nonparametric Friedman test was performed to determine whether a neuron’s responses at 12 orientations were significantly different from each other. To reduce Type-I errors, the significance level was set at α = 0.01. Third, for those showing significant orientation differences, the trial-based orientation responses of each neuron were fitted with a Gaussian model:

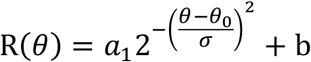

where R (*θ*) was the response at orientation *θ*, and the free parameters a_1_, *θ*_0_, *σ*, and b were the amplitude, peak orientation, standard deviation of the Gaussian function, and minimal response of the neuron, respectively.

More detailed information regarding the materials and methods can be found inJu et al. (2021), Ju et al. (2022), andLi et al. (2017).

### The transformer-based model

#### Input and output

To reconstruct a specific stimulus image seen by a monkey, the model input was a vector of neuronal responses whose length corresponded to the number of neurons. The model output was the reconstructed Gabor stimulus image of 79 × 79 pixels.

A total of 144 images were used, defined by a combination of twelve orientations, four SFs (1, 2, 4, and 8 cpd), and three sizes (s of the Gaussian envelope of the Gabor image).

Additional data with two lower SFs (0.25 & 0.5 cpd) were excluded as neurons tuned to these SFs were rare(Guan et al., 2021). To prevent overfitting, 10% of the conditions were randomly allocated as a validation set, which was not seen during training. Model training stopped when the accuracy on the validation set no longer increased, and this accuracy was taken as the best validation performance of the final model. The final dataset for further analysis consisted of 12 orientations at the preferred spatial frequency and size. For each stimulus condition, the input vectors were obtained by averaging the responses of twelve repeated trials, which enhanced the signal-to-noise ratio of the model inputs. In addition, these responses were normalized to the range [0, 1].

#### Embedding

The response vector *R*^0^ was first fed into the embedding module, which was associated with a distinct weight vector:

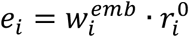

where *e*_*i*_ represented the embedded response of the *i*^*th*^ neuron, *w*_*i*_ was the independent weight assigned to the *i*^*th*^ neuron, and 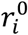 was the original response of the *i*^*th*^ neuron. Subsequently, the embedded responses for all neurons were concatenated to form the embedding matrix *R*^1^:

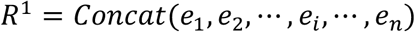

The embedding matrix *R*^1^ was obtained by applying an embedding transformation to the neuron responses. Its dimension was *n* × *d*_*model*_, where *n* was the number of neurons, and *d*_*model*_ was the dimensionality of the embedding for each neuron.

#### Positional encoding

Following the embedding process, positional encoding provided the sequence information:

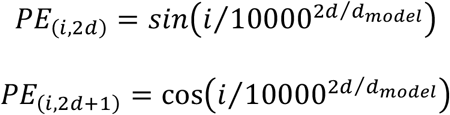

where *i* represented the position of a neuron in the sequence, and *d* denoted its position within the *d*_*model*_-dimensional embedding space. This approach to positional encoding generated a matrix of dimension *n* × *d*_*model*_. Here neurons were ordered according to their orientation preferences in the sequence, and neurons with close orientation preferences in the actual orientation space (e.g., 0° and 179°) received similar encodings due to the periodic nature of the sine and cosine functions. The dimension congruence allowed for the direct addition of the positional encoding to the embedding matrix, integrating position-based information into the neural response embeddings. The addition was performed elementwise:

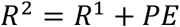

#### Self-attention

After the vector was enriched with response characteristics from the embedding and positional encoding stage, it was fed into the self-attention mechanism. This pivotal module computed attention scores by transforming the vector into queries (Q), keys (K), and values (V) using learned weight matrices:

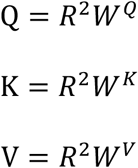

where *W*^*Q*^, *W*^*K*^, *W*^*V*^ were the weight matrices. Consequently, each neuron’s response was projected into a set of query, key, and value vectors, facilitating the calculation of attention scores.

For each neuron, a query (*Q*) was matched against all keys (*K*) to compute an attention map via a dot-product operation, which were then scaled, normalized, and passed through a SoftMax function to yield a probabilistic distribution of attention scores across the neurons:

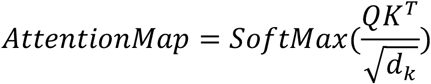

where *d*_*k*_ represented the dimensionality of the key vectors, which scaled the dot-product to control the variance of the attention scores.

The output of the *SoftMax*(*s*_*qk*_), for a given neuron’s query vector (indexed by *q*), was computed across all key vectors (indexed by *k*) and represented the attention distribution over all neurons:

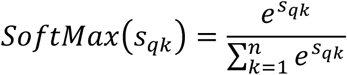

where *s*_*qk*_ was the *AttentionMap* between the *q*^*th*^ query and *k*^*th*^ key, and *n* was the total number of neurons. Subsequently, the attention scores were used to generate a weighted sum of the value vectors, producing a functional connection representation matrix for each neuron. The operation was mathematically expressed as:

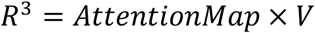

where *R*^3^ denoted the enriched per-neuron responses after self-attention was applied, *AttentionMap* represented the matrix of attention scores, and *V* was the matrix of value vectors.

#### Unembedding

At this stage, each neuron’s high-dimensional representation, encoded with interaction information from the self-attention mechanism, was projected back to a one-dimensional space. This reduction was conducted through a distinct set of learned weights assigned to each neuron:

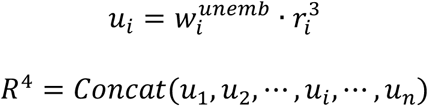

where *u*_*i*_ represented the unembedded response of the *i*^*th*^ neuron, 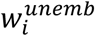 was the independent weight assigned to the *i*^*th*^ neuron, and 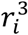 was the *i*^*th*^ element of *R*^3^.

The one-dimensional outputs could be directly compared to the original neural responses or used to reconstruct the visual stimulus, providing an endpoint for the model’s processing pipeline.

#### The feedforward layer

This layer processed the one-dimensional output vector from the unembedding stage, generating a reconstruction vector for each input. The transformation was guided by:

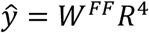

where ŷ was the vector that encapsulated the model’s prediction of the Gabor stimulus. Following the generation of the reconstruction vector, a reshaping operation was employed to transform these vectors into a two-dimensional image format.

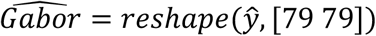

### Model training

During the training phase, the model aimed to minimize a loss function that quantified the discrepancy between the predicted and actual Gabor stimuli.

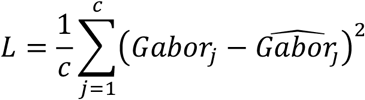

The loss function *L* was computed as the mean squared error (*MSE*) across conditions, where *Gobor*_*j*_ represented the actual image for the *j*^*th*^ condition, and 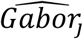 was the corresponding predicted image produced by the model. The objective of the training was to adjust the model parameters to minimize *L* across all conditions, so that the model was trained to predict the Gabor stimulus as accurately as possible, reducing the prediction error over successive training iterations. The model parameters to be trained are detailed in Table 1.

**Table 1.** Model parameters.

| Component | Parameters | Dimension |
| --- | --- | --- |
| Embedding | $W^{emb} = \text{Concat}(w_1^{emb}, w_2^{emb}, \dots, w_i^{emb}, \dots, w_n^{emb})$ | $[n \times d_{model}]$ |
| Self-attention | $W^Q \ W^K \ W^V$ | $[d_{model} \times d_{model}] \times 3$ |
| Unembedding | $W^{unemb} = \text{Concat}(w_1^{unemb}, w_2^{unemb}, \dots, w_i^{unemb}, \dots, w_n^{unemb})$ | $[n \times d_{model}]$ |
| Feedforward | $W^{FF}$ | $[n \times 79^2]$ |

We used the RMSprop optimizer to optimize model parameters. RMSprop is an adaptive learning rate method, which enhanced convergence and minimized oscillations during training by adjusting the learning rate based on the moving average of squared gradients. This ensured stability throughout the training process. The algorithm proceeded as follows:

First, the exponentially weighted moving average of the squared gradients was computed as:

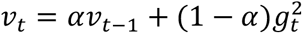

where *v*_*t*_ was the moving average of the squared gradients, *α* was the smoothing factor set to 0.9 in our experiments, and *g*_*t*_ was the current gradient.

Next, the parameters were updated as:

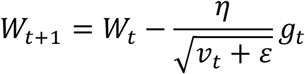

where *W*_*t*_ represented the model parameters at step *t, η* was the learning rate set to 0.00005, *v*_*t*_ was the moving average of the squared gradients, and *ε* was a small constant to prevent division by zero.

### Self-attention modulation

To account for the influence of self-attention on neuronal responses, a modulation formula was employed to adjust the original response *R*_*j*_ of each neuron under the *j*^*th*^ condition based on its attention score. The modulated response 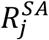 for each neuron was computed as follows:

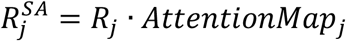

where *R*_*j*_ represented the neuron’s original response under the *j*^*th*^ condition, and *Attentation Map*_*j*_ was the attention score derived from the self-attention mechanism’s assessment of inter-neuronal influences under the same condition. 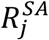 was the response of the neuron after modulation by the self-attention mechanism.

The attention map served as a dynamic filter that selectively amplified or attenuated neuronal responses based on their relevance to the task, enhancing signal processing efficiency and focusing attention on the most pertinent information.

### Structural similarity index measure (SSIM)

The SSIM between two images *x* and *y* of common size 79 × 79 was defined as:

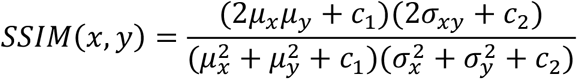

where *μ*_*x*_ and *μ*_*y*_ were the average intensities of *x* and *y*, 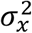 and 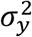 were the variances of *x* and *y, σ*_*xy*_ was the covariance between *x* and *y*, and *c*_1_ and *c*_2_ were constants to stabilize the division with weak denominator.

### Gabor orientation extraction

Let *I*(*x, y*) represent the Gabor image. The horizontal (*G*_*x*_) and vertical (*G*_*y*_) gradients of the image were computed as:

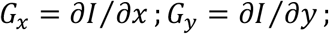

The gradients were then combined to form a matrix of gradient vectors. For an image with 6241 (79^2^) pixels, the gradient vectors were represented as:

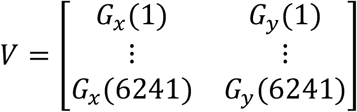

The principal component analysis (PCA) was then applied to the matrix *V*, resulting in the coefficient matrix *coeff*:

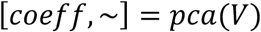

where, *coeff* was the matrix of eigenvectors (principal components), in which each column represented a principal component direction. The first principal component *coeff*(: ,1) corresponded to the direction of maximum variance in the gradient data.

The primary direction *θ* was calculated using the arctangent function, defined as:

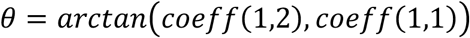

where *coeff*(1,2) and *coeff*(1,1) were the elements of the principal component vector obtained from PCA.

The angle *θ* in radians was converted to degree using the following formula:

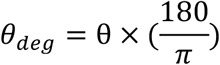

The orientation was then adjusted to be within the range [0, 180):

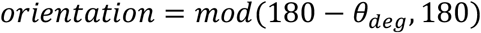

Following these steps, the orientation information in the Gabor image was effectively extracted, providing a reliable method for determining the dominant orientation of image features.

### Orientation gradient

To quantify the local variation of neuronal orientation preferences, a gradient map was calculated for the orientation dataset. For each neuron, neurons within 30 μm in cortical space were selected as its local neighborhood. The orientation difference between the neuron and each neighboring neuron was computed as the minimal angular difference:

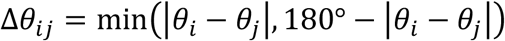

This angular difference was then normalized by the cortical distance between the two neurons:

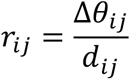

where *d*_*ij*_ represents the Euclidean distance between neurons *i* and *j*. The local orientation gradient value of the neuron was obtained by averaging all *r*_*ij*_ values within its neighborhood:

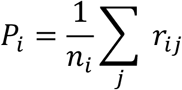

where *n*_*i*_ is the number of neighboring neurons within 30 μm. Neurons without any neighbors within this range were assigned a gradient value of zero.

The discrete orientation gradient values of all neurons were then mapped back into the image space by assigning each neuron’s gradient value to its corresponding pixels. To generate a continuous orientation gradient map, the discrete map was smoothed using a Gaussian filter with a standard deviation of *σ* = 15 pixels:

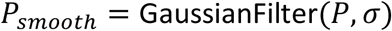

### Noise correlation

To quantify trial-to-trial response co-variability, noise correlations were calculated between all pairs of neurons for each stimulus orientation. For each neuron, responses at its preferred spatial frequency and size were extracted, with each orientation presented 12 times. The mean response across 12 trials was subtracted to obtain residuals, which were then z-scored to normalize variability across neurons, following the procedure described byChurchland et al. (2010). Pairwise Pearson correlation coefficients were computed between the z-scored residuals of all neuron pairs. This process was repeated for all stimulus orientations.

### Control decoding models

We implemented two commonly used neural decoding models as controls: a linear decoder and a nonlinear multilayer perceptron (MLP). Because orientation is a circular variable with 180^º^ periodicity, all decoders predicted orientation using a two-dimensional circular representation rather than directly regressing angular values. Specifically, each orientation label was encoded as:

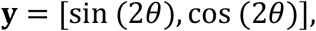

where *θ* denotes the stimulus orientation. This double-angle representation maps orientations separated by 180° onto the same point while preserving the relative angular relationship between orientations. The decoder output consisted of two continuous values:

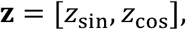

corresponding to the predicted sine and cosine components. The output vector was normalized before calculating the loss:

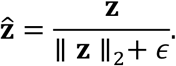

The predicted orientation was recovered using the inverse transformation:

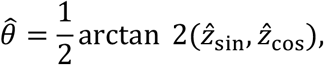

And converted into degrees within the range of 0° to 180°. All decoders were optimized using a cosine-based circular loss: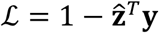, where 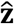 and **y** denote the normalized predicted and target orientation vectors, respectively.

The first control model was a linear decoder, consisting of a single fully-connected layer that directly mapped the population neural response **r** to the two-dimensional orientation representation: **z** = **W**_*linear*_ **r + b**_*linear*_. This model contained no hidden layer or nonlinear activation and therefore provided a baseline for orientation information recoverable through fixed linear weighting of neuronal activity.

The second control model was a nonlinear multilayer perceptron containing a single hidden layer and nonlinear activation. Specifically, the hidden representation was calculated as:

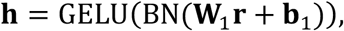

where BN denotes batch normalization and GELU denotes the nonlinear activation function. The hidden layer contained 128 units. The hidden representation was subsequently mapped to the orientation output space: **z** = **W**_2_ **h + b**_2_.

These two models allowed us to distinguish the contribution of fixed linear pooling and generic nonlinear feedforward computation from the attention-based population readout implemented by the transformer. Both control models were trained using the AdamW optimizer with an initial learning rate of 0.001 and a weight decay coefficient of 0.0001. The learning rate was reduced by a factor of 0.5 when the evaluation loss did not improve for 100 epochs. Training was performed for a maximum of 5,000 epochs with early stopping when the evaluation loss failed to improve for 1,000 consecutive epochs. The batch size was set to 16 for both control models. All control models used the same circular orientation loss and optimization strategy.

## Acknowledgments

This study was supported by a National Natural Science Foundation of China, Brain Science and Brain-like Intelligence Technology -National Science and Technology Major Project grant (2022ZD0204600) to SMT and CY.

